# Thermodynamic modelling of synthetic communities predicts minimum free energy requirements for sulfate reduction and methanogenesis

**DOI:** 10.1101/857276

**Authors:** Hadrien Delattre, Jing Chen, Matthew Wade, Orkun S Soyer

## Abstract

Microbial communities are complex dynamical systems harbouring many species interacting together to implement higher-level functions. Among these higher-level functions, conversion of organic matter into simpler building blocks by microbial communities underpins biogeochemical cycles and animal and plant nutrition, and is exploited in biotechnology. A prerequisite to predicting the dynamics and stability of community-mediated metabolic conversions, is the development and calibration of appropriate mathematical models. Here, we present a generic, extendable thermodynamic model for community dynamics accounting explicitly for metabolic activities of composing microbes, system pH, and chemical exchanges. We calibrate a key parameter of this thermodynamic model, the minimum energy requirement associated with growth-supporting metabolic pathways, using experimental population dynamics data from synthetic communities composed of a sulfate reducer and two methanogens. Our findings show that accounting for thermodynamics is necessary in capturing experimental population dynamics of these synthetic communities that feature relevant species utilising low-energy growth pathways. Furthermore, they provide the first estimates for minimum energy requirements of methanogenesis and elaborates on previous estimates of lactate fermentation by sulfate reducers. The open-source nature of the developed model and demonstration of its use for estimating a key thermodynamic parameter should facilitate further thermodynamic modelling of microbial communities.

## INTRODUCTION

Microbial communities are found in diverse habitats including the oceans, soil, animal guts, and plant roots. The interconnected metabolic activities in these microbial communities underpin the biogeochemical cycles that feed into the Earth’s ecosystem [1] and the nutrient cycles that support the growth of animals and plants [2, 3]. The same community level metabolic activities, and in particular anaerobic digestion (AD), are also exploited in biotechnology for water treatment and bioenergy production from organic waste [4]. Thus, the ability to capture microbial growth rates and metabolic activities within microbial communities is identified as an important prerequisite for the predictive modelling of planetary ecosystem dynamics, animal and plant health, and biotechnological waste valorisation [4].

Modelling microbial community dynamics is a significant challenge due to the complexity of these systems. Typical communities, for example those found in human gut or AD reactors, are composed of 100s to 1000s of distinct microbial species [5, 6]. The metabolic activities, and hence the growth, of these different species are interlinked to each other through metabolic interactions that resemble ecological ones [7]. This resemblance has motivated the adaptation of simplified ecological models (e.g. Lotka-Volterra models) to the modelling of microbial communities [8]. While these models allow drawing generalised hypotheses about the role of different types of interactions on microbial community stability [9], they do not capture metabolite dynamics, which are shown to be essential for predicting population dynamics [10].

Microbial growth and metabolite dynamics are historically captured by empirical models such as the Monod growth function [11, 12]. These growth functions have been used to capture dynamics of microbial communities, most notably to construct relatively large-scale models describing AD communities, as used in wastewater treatment engineering [13]. These models reduce system complexity by considering functional groups (so-called ‘guilds’), rather than individual species, thereby capturing key metabolic processes and interactions such as polymer degradation, sulfate reduction, and methanogenesis [14, 15]. The guild-based approach makes it possible to calibrate and test these models against the key metabolites measured in AD reactors, bringing us closer to predicting performance and stability of communities in biotechnological applications. Towards achieving this goal, however, a key limitation has been the inadequacy of Monod-type models to capture microbial metabolic conversions that are at a low energy level, and thus operating close to thermodynamic equilibrium [16, 17]. Such ‘thermodynamic inhibition’ of microbial growth and metabolism is highly relevant to AD, as well as soil, sediment, and gut communities, where there is commonly a depletion of strong electron acceptors and a shift of metabolism from high energy respiratory pathways to low energy fermentative pathways [18].

To capture thermodynamic inhibition effects, a simple thermodynamic model has been proposed that adjusts a Monod-type growth function with a thermodynamic factor based on the free energy of the growth-supporting metabolic conversion [16, 17, 19, 20]. This approach is further elaborated upon by considering the fact that part of the free energy from a given metabolic conversion must be invested into cellular maintenance and as a metabolic driving force, thus defining a minimal energy threshold for a growth-supporting pathway [21]. Incorporating such a thermodynamic model has allowed studying the basis of observed diversity in microbial communities [22] and making qualitative predictions on population dynamics in microbial communities [23]. A fully quantitative prediction of population and metabolite dynamics, however, requires that these models implement specific, calibrated kinetic and thermodynamic parameters for each of the accounted microbial species and their metabolic conversions.

Kinetic parameters of microbial growth have been collected over decades of research using monocultures grown under defined conditions. In particular, maximal growth rate (*v*_*max*_), substrate affinity coefficient (*K*_*s*_), and biomass yield from substrate (*Y*_*s/x*_) have been experimentally estimated for individual species that represent common functional groups seen in microbial communities. For some of these kinetic parameters, in particular biomass yield and substrate uptake rate, calibrated methods have been derived that can predict parameters from existing data and first principles approximations [24–26]. The key thermodynamic parameter, namely the minimum energy threshold of different metabolic conversions, however, remains mostly unavailable. Moreover, there has not been any focussed exploration of what kind of experimental measurements can provide sufficiently robust estimations for this parameter. This situation limits the applicability of thermodynamic community models, which are required to fully capture the growth of many functionally relevant species.

Here, we aim to address this gap and develop a generic, readily extendable thermodynamic community model and use it to estimate the minimal free energy parameters from experimental time-series data. The model implements the multiple and distinct growth-supporting metabolic conversions possible in each organism and accounts for their possible thermodynamic limitations. It also accounts for metabolite phase exchanges and system pH. We calibrate the model using experimental data from synthetic communities composed of microbial species that represent key functional groups in AD systems; *Desulfovibrio vulgaris* (*Dv*), a sulfate reducer, *Methanococcus maripaludis* (*Mm*), a hydrogenotrophic methanogen, and *Methanosarcina barkeri* (*Mb*), a methanogen capable of acetoclastic methanogenesis. Using daily metabolite measurement from mono-, co-, and tri-cultures over a 21-day experiment, we show that the resulting model provides a superior fit to data, compared to a non-thermodynamic model, and that some of the thermodynamic parameters of the model can be calibrated using time-series data. These results show that thermodynamic models are appropriate and are needed to accurately capture metabolite dynamics in microbial communities, but that their full calibration requires a greater breadth of experimental data.

## MATERIALS AND METHODS

### Overall model description and availability

The model presented here aims to capture the population and metabolic dynamics within a microbial monoculture or a multi-species community. The model accounts for a set of growth-supporting metabolic pathways that involve either specific metabolites or cellular biomass (Figure 1). The chemical speciation of metabolites, as well as their exchange between gas and liquid phases is accounted for. The medium pH is also simulated based on the set of acid-base reactions that are included. The model is developed in a generic and user-accessible manner, so that growth-supporting reactions, species involved, gas/liquid exchange reactions, and acid-base reactions can be supplied by the user without any pre-requisite programming skills, and the source-code is extendable by advanced users. The entire model is encoded in an object-oriented software using Python 3.4 and the source-code and simulation manual are provided via authors’ research website at https://github.com/OSS-Lab/micodymora.

**Figure 1.**
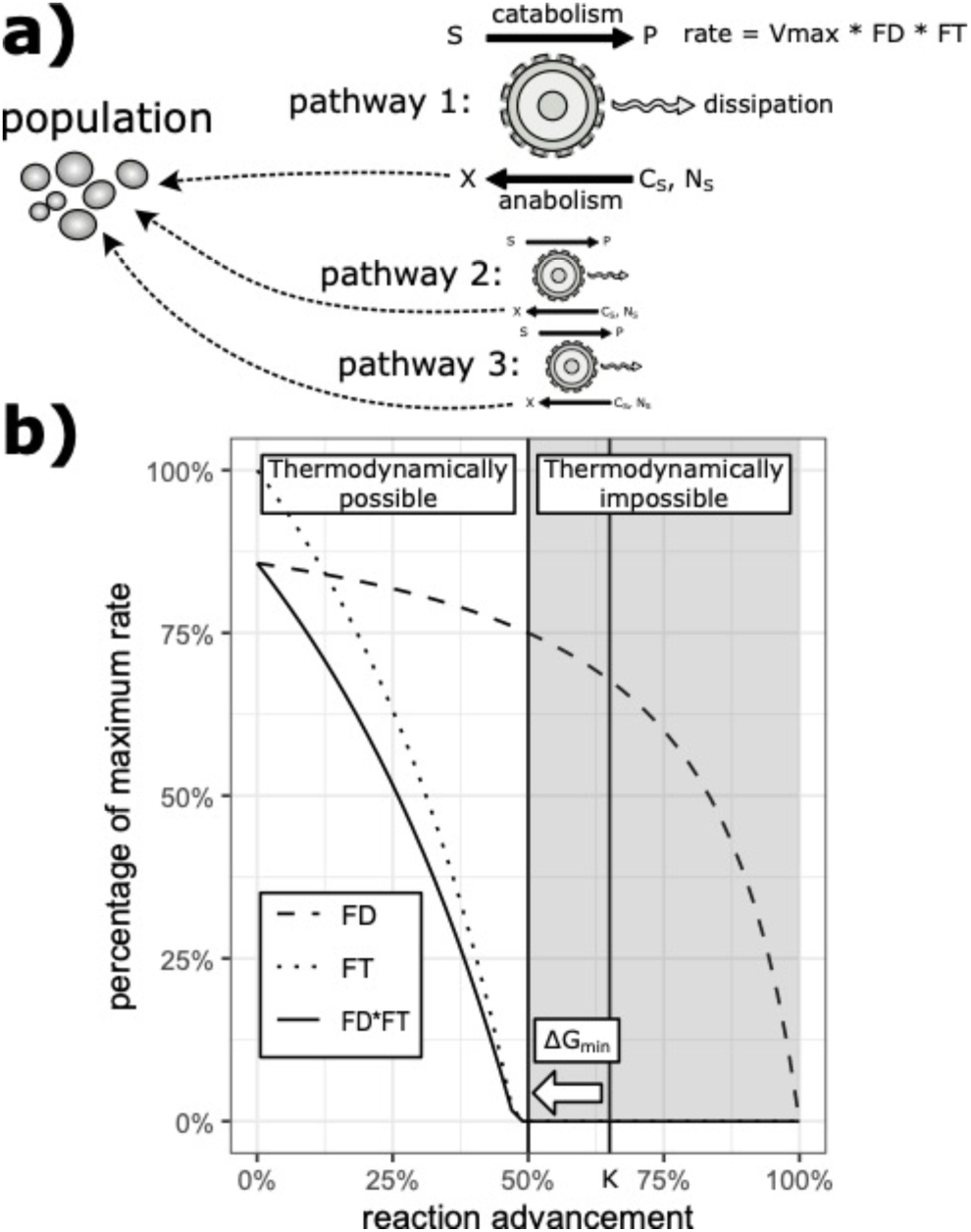
(**A)** Graphical summary of the presented model. Each microbial population is able to catalyse one to multiple different metabolic pathways. Each pathway consists in a catabolic reaction energetically coupled to an anabolic reaction. The rate of the catabolic reaction is given by a constant *v*_*max*_ multiplied by the *F*_*D*_ and *F*_*T*_ factors, representing enzyme kinetics and thermodynamics constraints respectively (see *Materials and Methods* and Equation 2). **(B)** Cartoon representation of the value of the *F*_*D*_ and *F*_*T*_ factors as a function of reaction advancement (this representation assumes a simple one-to-one substrate to product stoichiometry). Note that the greater the *ΔG*_*min*_ parameter is (as a negative number), the further left the point where *FD* becomes null moves.

### Growth supporting metabolic pathways

For the presented model, several growth-supporting (i.e. catabolic) and biomass forming (i.e. anabolic) metabolic pathways are considered to be encoded by *Dv*, *Mm* and *Mb* populations, as illustrated in Figure 1A and listed in Table 1. The anabolic (biomass producing) reactions of *Dv*, *Mm* and *Mb* populations are considered to utilise lactate, carbon dioxide, and acetate respectively as carbon source. In each of these reactions, biomass is represented as a generic molecule (with chemical formula C_1_H_1.8_O_0.5_N_0.2_), having an associated Gibbs free energy of formation of −67 kJ/mol [27]. Table 1 lists all growth-supporting catabolic reactions modelled in this work, along with their associated Gibbs free energy (*ΔG*^0^’) calculated at a pH of 7.0 and a temperature of 310.15 K with reagents other than protons in their standard state.

**Table 1:** Catabolic and anabolic reactions encoded by each species simulated in the presented model. Reaction Gibbs free energies are also shown, as calculated for pH=7, 1atm, and 310.15 K.

| Organism | Catabolic pathway(s) | Anabolic pathway | Dissipated energy |
| --- | --- | --- | --- |
| *Dv* | $\text{C}_3\text{H}_5\text{O}_3^- + 2 \text{H}_2\text{O} \rightarrow \text{C}_2\text{H}_3\text{O}_2^- + 2 \text{H}_2(\text{aq}) + \text{HCO}_3^- + \text{H}^+$ $\Delta G^{0'} = 27.58 \text{ kJ/mol}$ | $0.35 \text{C}_3\text{H}_5\text{O}_3^- + 0.2 \text{NH}_4^+ + 0.1 \text{H}^+ \rightarrow \text{C}_1\text{H}_{1.8}\text{O}_{0.5}\text{N}_{0.2} + 0.05 \text{C}_2\text{H}_5\text{O}_2^- + 0.4 \text{H}_2\text{O}$ $\Delta G^{0'} = 18.11 \text{ kJ/mol}$ | $\Delta G_{\text{dis}} = -265.1 \text{ kJ/mol}$ |
| | $\text{C}_3\text{H}_5\text{O}_3^- + 0.5 \text{SO}_4^{2-} \rightarrow \text{C}_2\text{H}_5\text{O}_2^- + \text{HCO}_3^- + 0.5 \text{H}_2\text{S}(\text{aq})$ $\Delta G^{0'} = -170.17 \text{ kJ/mol}$ | | $\Delta G_{\text{dis}} = -228.2 \text{ kJ/mol}$ |
| | $\text{H}_2(\text{aq}) + 0.25 \text{SO}_4^{2-} + 0.5 \text{H}^+ \rightarrow 0.25 \text{H}_2\text{S}(\text{aq}) + \text{H}_2\text{O}$<br>$\Delta G^{0'} = -56.33 \text{ kJ/mol}$ | | $\Delta G_{\text{dis}} = -362.1 \text{ kJ/mol}$ |
| *Mm* | $0.25 \text{HCO}_3^- + \text{H}_2(\text{aq}) + 0.25 \text{H}^+ \rightarrow 0.25 \text{CH}_4(\text{aq}) + 0.75 \text{H}_2\text{O}$<br>$\Delta G^{0'} = -48.12 \text{ kJ/mol}$ | $\text{HCO}_3^- + 2.1 \text{H}_2(\text{aq}) + 0.2 \text{NH}_4^+ + 0.8 \text{H}^+ \rightarrow \text{C}_1\text{H}_{1.8}\text{O}_{0.5}\text{N}_{0.2} + 2.5 \text{H}_2\text{O}$<br>$\Delta G^{0'} = -64.73 \text{ kJ/mol}$ | $\Delta G_{\text{dis}} = -876.4 \text{ kJ/mol}$ |
| *Mb* | $0.25 \text{HCO}_3^- + \text{H}_2(\text{aq}) + 0.25 \text{H}^+ \rightarrow 0.25 \text{CH}_4(\text{aq}) + 0.75 \text{H}_2\text{O}$<br>$\Delta G^{0'} = -48.12 \text{ kJ/mol}$<br>$\text{C}_2\text{H}_3\text{O}_2^- + \text{H}_2\text{O} \rightarrow \text{CH}_4(\text{aq}) + \text{HCO}_3^-$<br>$\Delta G^{0'} = -14.65 \text{ kJ/mol}$ | $0.525 \text{C}_2\text{H}_3\text{O}_2^- + 0.2 \text{NH}_4^+ + 0.275 \text{H}^+ \rightarrow \text{C}_1\text{H}_{1.8}\text{O}_{0.5}\text{N}_{0.2} + 0.05 \text{HCO}_3^- + 0.4 \text{H}_2\text{O}$<br>$\Delta G^{0'} = 28.63 \text{ kJ/mol}$ | $\Delta G_{\text{dis}} = -1059.8 \text{ kJ/mol}$<br>$\Delta G_{\text{dis}} = -1524.0 \text{ kJ/mol}$ |

### Modelling population and metabolite dynamics

To model population dynamics, we consider the *Dv*, *Mm* and *Mb* populations as implementing catabolic pathways available for each species. The overall dynamics of each population are governed by a differential equation that accounts for all of its catabolic pathways rates, as well as its anabolic biomass production. It is assumed that for a given population, the anabolic reaction has the same formula for each of its catabolic pathways (see Table 1). To represent the moles of biomass formation per moles of catabolite consumed, the stoichiometry of each anabolic reaction is multiplied by a dynamic yield coefficient (*Y*, in mol_X_/mol_S_ where X stands for biomass and S for the substrate by which the catabolic pathway’s formula is normalized). This yield coefficient is computed dynamically from the energetics of catabolic and anabolic reactions, as well as the dissipated energy during biomass formation;

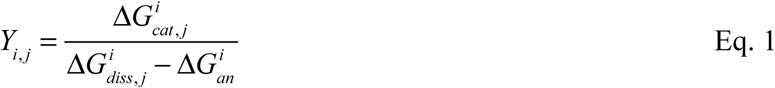

where the indices *i* and *j* range over a given species (e.g. *Dv*) and catabolic pathway (e.g. lactate fermentation) respectively. The 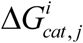 is the Gibbs free energy (in kJ/molS), of the associated catabolic pathway *j* in a given species *i*, while 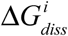 is the dissipated energy in anabolism for the same pathway. 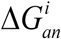 is the Gibbs free energy of the anabolic reaction respectively (both in kJ/mol_X_) for species *i*. The Gibbs free energy of the catabolic and anabolic reactions are calculated dynamically during model simulation from the chemical species concentrations. The dissipated energy is that which is harvested by the population through its catabolism and which is not chemically stored as biomass. It encompasses a wide diversity of processes including heat and entropy emission and cellular maintenance. Its value is estimated based on experimentally measured dissipated energy for different microbial species [28] (see Table 1).

The specific rate (*r*_*i,j*_ in mol_S_/(mol_X_∙hour)) of a catabolic pathway *j* for a given species *i* is given by;

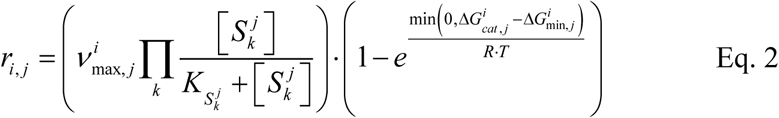

where 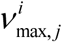 is the maximum catabolic turnover rate, expressed in mol_S_/(mol_X_∙hour) and specific to the pathway *j* and the population *i*, 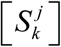 is the concentration of the *k*^th^ limiting substrate of the pathway *j*, 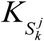 is the half-saturation coefficient for that substrate, 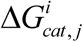 is the Gibbs free energy of the catabolic reaction *j* for species *i*, and 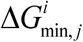 is the minimum energy threshold for that catabolic reaction. This last term 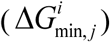 captures how energy-storing reactions coupled to the catabolic reaction (e.g. conserved moieties regeneration, proton extrusion, etc.) affects its rate. *R* (in J/(mol∙K)) and *T* (in K) denote the gas constant and system temperature respectively. Note that in the main text, we refer to the first and second terms of Equation 2 as kinetic (*F*_*D*_) and thermodynamic (*F*_*T*_) factors, similar to previous presentations in the literature [29]. The kinetic coefficients 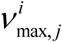 and 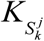 are compiled from the literature and are listed in Supplementary Table S1.

Using the rate of the catabolic pathway and biomass yield of the anabolic pathway, we can write a differential equation describing the dynamics of the concentration of any chemical *A* according to the catalytic activity of each population;

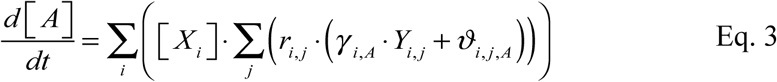

where [*X*_*i*_] represent the biomass concentration of the *i*’th population, *γ*_*i, A*_ is the stoichiometric coefficient for chemical *A* in the anabolic reaction of the *i*’th population, and *ϑ*_*i,j,A*_ is the stoichiometric coefficient for chemical *A* in the *j*’th catabolic pathway of the *i*’th population.

The dynamics of the biomass associated to a population *i* obeys essentially the same equation as for the concentration of chemical (Equation 3), however, *ϑ*_*i,j,X*_ is always zero because biomass is not produced or consumed by catabolism and *γ*_*i,X*_ is always one because the anabolic formula is normalized to the mole of biomass. Additionally, we account for the loss of biomass through death using a linear decay coefficient *k*_*d*_ (1/hour), resulting in the following differential equation for biomass;

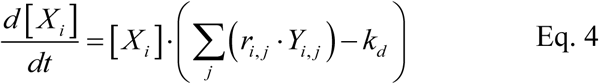

We assume in the current model that *k*_*d*_ (=8.33e-4 1/h) is the same for all species. This value is based on the experimental estimates made on *Desulfovibrio vulgaris* monocultures [30].

### Modelling of gas/liquid transfer

Some chemicals exist in both the gas and liquid phases. Any of such chemical species, *A*, is accounted for as two separate chemical species *A(aq)* and *A(g)* respectively. The concentrations of each species are accounted for by moles per liter of their respective phase volume. The transfer dynamics occurring between the two forms is captured through a set of differential equations given by;

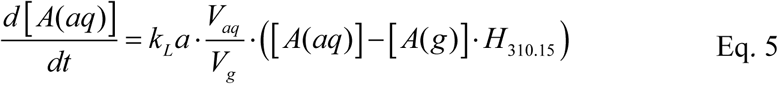

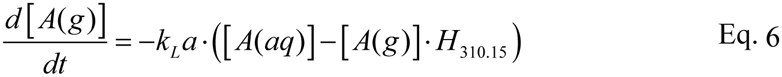

where *k*_*L*_*a* is the mass transfer coefficient of the chemical (in 1/hour) [31], *H*_*310.15*_ is the Henry constant of the chemical at 310.15K (and expressed in mol/(m^3^∙Pa)), *V*_*aq*_ is the volume of the liquid phase and *V*_*g*_ is the volume of the gas phase (both in liters). Henry constants were obtained from the literature [32], and adjusted for a temperature of 310.15 K using the relation between Henry’s constant and the solution enthalpy (Δ*H*_*sol*_) as follows; 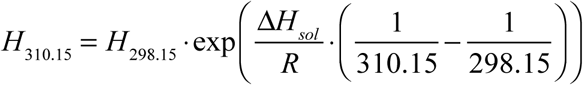. The list of species that are modelled as distributed between liquid and gas phases, and their associated Henry constants and mass transfer coefficients are listed in Supplementary Table S2.

### Modelling of medium pH

At the beginning of each timestep in the integration of the differential equation system (composed of Equations 3–6), the pH of the solution is determined. This is done by solving the charge balance of the system using the Brent method [33], while considering the proportion of each ionized species depending on the pH. The acid-base equilibria that are considered and determined at each time step are listed in Supplementary Table S3, along with the associated pK values.

### Model parameters and parameter calibration

The kinetic parameters used in the model are listed in Table S1, and are based on experimental estimates given in the literature. Henry constant and mass transfer coefficients (Table S2) are compiled from the literature or measured in this study (see below). The thermodynamic parameters are either calculated dynamically (as explained above) or adapted from the literature (Table 1). The only parameters of the model that are calibrated against experimental data are the 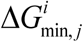 of the different catabolic pathways. These parameters are calibrated using a recently introduced optimisation procedure [34]. This approach has been chosen because it has specifically been proposed in the context of the estimating microbial growth parameters and to circumvent the problem of parameter identifiability [35]. In brief, this approach calibrates multiple parameters (the 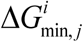 of each pathway) against multiple observed variables (experimentally observed lactate, acetate, H_2_(g) and CH_4_(g) concentrations). With this optimisation procedure, the parameters are treated in a hierarchical fashion according to two properties; the extent (the number of variables affected upon changing a given parameter) and scale (the level of change in variables induced by changing a given parameter) of their effect. The parameter that produces the strongest effect amongst the least number of variables is selected first and the experimental time course data of the variables are weighted so that those variables that are most affected by the parameter have more weight in the calculation of the match between model prediction and experimental data during the optimisation procedure. The selected parameter is then optimized against the weighted experimental data on variables using the truncated Newton method (here we use the implementation available in the Python 3.4, package “scipy”). This method minimizes the weighted sum of squared distance between the model predictions and the experimental data on the variables. Once a parameter is optimized in this way, its value is fixed and removed from the list of the parameters to be optimized. The optimisation procedure then restarts with the remaining parameters until they have all been optimized and then repeated again for a different set of starting values. The whole process of optimisation is repeated until it yields no significant improvement anymore in terms of distance between the model predictions and the experimental data on variables.

### Numerical simulations

The model is used to simulate the dynamics of the different populations as well as the key metabolites and system pH. Simulations were run to emulate the actual experiments in terms of run duration and initial starting conditions. The latter was assumed to be an equal biomass distribution among constituting species. To estimate this distribution, total biomass concentration in C-mol/L was approximated from the experimental OD (at 600nm) measurements at the start of the experiment (Supplementary Table S4) and using a previously calibrated relationship between OD and biomass using sulfate reducing bacteria (predominantly *Desulfovibrio vulgaris*) [36]; ln(*DW*) = 5.12 ∙ OD600 – 4.987, where *DW* is the dry weight of the cells in g/L. We converted the resulting *DW* value to 1/C-mol by dividing it by the molecular weight of the generic molecule used to represent biomass (C_1_H_1.8_O_0.5_N_0.2_, [27]); 24.6 g.C/mol. The resulting biomass concentration was then evenly distributed between the existing populations to create the initial point for simulations.

### Experimental estimation of *k*_*L*_*a* for H_2_, CO_2_ and CH_4_

The *k*_*L*_*a* parameter for H_2_, CO_2_ and CH_4_ was estimated based on experimental measurements using the same setup as in our experimental system. Anaerobic medium was prepared as previously described [37], containing 30mM Na-lactate and 7.5mM Na_2_SO_4_. Anaerobic culture tubes (Hungate tubes, Chemglass Life Sciences, Vine-land, NJ, USA) were prepared with 5mL medium and 0.1mL of 100 mM Na_2_S•9H_2_O in each tube, sealed in an anaerobic chamber station (MG500, Don Whitley) and autoclaved. The headspace gas pressure and composition of the tubes were measured using a micro gas-chromatograph (GC) (Agilent 490 micro-GC, Agilent Technologies) and recorded. A gas mixture of H_2_, CO_2_ and CH_4_ was prepared by first flushing two 118mL serum bottles with 80% H_2_ / 20% CO_2_ gas mixture for 3 minutes at 0.5 L/min flow rate and balancing the final pressure to 1atm (101325 Pa). Then, 10mL of 90% CH_4_ / 10% CO_2_ gas mixture at 1atm was injected into each serum bottle using a gas tight glass syringe (Cadence Science, Inc., Italy). 2.0mL of the resulting gas mixture is injected into each of the prepared Hungate tubes using a gas tight glass syringe. The tubes were incubated under 37°C for more than 24 hours, in order to let the added gas to be equilibrated between the headspace and the aqueous phase. The tubes were then flushed with 100% N_2_ for 2 minutes at a flow rate of 0.2 L/min and their headspace pressure brought to 1atm using sterile needle and filter. The tubes were then returned to the 37°C incubator, and brought out in replicates of 3 for temporal measurement of headspace gas composition at pre-determined intervals of 0, 1, 2, 4, 8 and 24 hours. The resulting temporal gas equilibration data is then used to estimate the *k*_*L*_*a* value for H_2_, CO_2_, and CH_4_. Specifically, the *k*_*L*_*a* values were obtained by minimizing the sum of squared error between average observed measurements and the integration of the dynamics of gas transfer considered (see Equations 5 & 6).

### Experimental implementation of monocultures and synthetic microbial communities

The three strains of *Dv*, *Mb* and *Mm* and anaerobic medium preparations were done as previously described [37]. In brief, the mono-cultures of *Dv*, *Mb* and *Mm* were cultivated in 5mL anaerobic media for 4, 21 and 7 days respectively to reach their late log phase. These monocultures were all grown at 37°C in the same anaerobic medium base (OSM1.0 media as described in [37]), but with different carbon and energy sources; 30mM Na-lactate and 10mM Na_2_SO_4_ for *Dv*, 100mM Na-acetate for *Mb* and 10mM Na-pyruvate and 68.4mM NaCl for *Mm*. For the last species, the headspace is also filled with 80%H_2_ / 20%CO_2_ gas mixture at a pressure of 2atm. To create synthetic communities of co- and tri-cultures, we first created stock cultures by taking 2mL aliquots of each monoculture using sterile needle syringe inside anaerobic chamber and inoculating these in the combinations of *Dv-Mb*, *Dv-Mm* and *Dv-Mb-Mm* into different serum bottles, which contained 50mL OSM1.0 medium with 30mM Na-lactate and 7.5mM Na_2_SO_4_. The inoculated serum bottles were placed in a 37°C incubator for 21 days. At the end of this period, 17.5mL cultures of different combinations from the incubated serum bottles were transferred into 500mL anaerobic Duran bottles containing 350mL of the above medium. The Duran bottles were linked to a Micro-GC (Agilent 490 micro-GC, Agilent Technologies) for the continuous monitoring of the methane production over two weeks. The active methanogenic communities in all combinations are confirmed in this way and the cultures were considered and used as the stock cultures for the following step. 5mL of the stock cultures from each combination were extracted inside anaerobic chamber, mixed separately with 5mL fresh anaerobic medium OSM1.0 with 30mM Na-lactate and 7.5mM Na_2_SO_4_ in Hungate tubes, and incubated at 37 °C for 7 days. These cultures formed the inocula for the following time-series experiment.

To measure temporal dynamics of co- and tri-cultures, as well as *Dv* monoculture, we designed a time-series experiment that involved starting a large number of replicate tubes and terminating a set of this large batch at different time points for gas and metabolite measurements. In total, 273 anaerobic Hungate tubes were prepared to collect data for 21 time points. Each tube contained 5mL OSM1.0 medium with 30mM Na-lactate and 7.5mM Na_2_SO_4_. According to the full reaction of sulfate reduction by *Dv* in Table 1, 7.5mM sulfate should allow *Dv* to convert 15mM lactate fully, while the conversion of the other 15mM lactate would rely on *Dv*’s other less thermodynamically favourable pathways. The tubes were numbered individually and separated into 21 batches. Each batch contains 13 tubes, of which 1 tube was used as blank control and 3 replicate tubes were used each for the 4 cultures: *Dv*-*Mb*, *Dv*-*Mm*, *Dv*-*Mb*-*Mm* and *Dv*, respectively. The tubes were inoculated with the respective cultures using the stock cultures described above, and following the tube and batch numbers. The initial optical density (OD) at 600nm and headspace pressure were recorded for each tube using a spectrophotometer (Spectronic 200E, Thermo Scientific) and a needle pressure gauge (ASHCROFT 310, USA). All tubes were incubated at 37°C. Over the following 21 days, 13 tubes of one batch were terminated on each single day to measure their OD at 600nm, pH (Mettler Toledo M300, Columbus, Ohio, USA), gas pressure (ASHCROFT 310, USA), gas composition using Micro-GC (Agilent 490 micro-GC, Agilent Technologies) and the lactate, acetate, pyruvate and sulfate concentrations using Ion Chromatography (Dionex ICS-5000^+^ DP, Thermo Scientific) as described previously [37].

## RESULTS AND DISCUSSION

To develop and calibrate a thermodynamic model of microbial growth and metabolite dynamics in a community context, we focus here on defined anaerobic synthetic communities. In particular, we use a recently developed experimental model system for studying syntrophic interactions among sulfate reducers and methanogens [37, 38], which make up a key part of anaerobic microbial communities found in AD reactors and freshwater and estuary sediments. The studied synthetic systems are composed of a representative sulfate reducer (*Desulfovibrio vulgaris*, *Dv*), and two different methanogens representing hydrogenotrophic (*Methanococcus maripaludis*, *Mm*) and hydrogeno/acetotrophic (*Methanosarcina barkeri*, *Mb*) methanogenesis pathways (see Materials and *Methods* and Figure 1). We have collected here data on metabolite dynamics over a 3-week period from *Dv* monocultures, *Dv*-*Mm* and *Dv*-*Mb* co-cultures, and *Dv*-*Mm*-*Mb* tri-cultures under specific media conditions (see *Materials and Methods*).

### A comprehensive and generic thermodynamic model of community dynamics

To capture community and metabolite dynamics, we developed a comprehensive and expandable thermodynamic model that also accounts for metabolite phase exchanges and medium pH (see *Materials and Methods*). As is common for many microbes found in microbial communities, the species composing the studied synthetic communities can catalyse multiple, distinct metabolic pathways, sometimes starting from the same substrate. To account for these different metabolic activities of each species, we considered that each population can utilise any number of pathways at once, as previously described [30]. Each pathway consists of a catabolic (energy harvesting) reaction and an anabolic (biomass synthesis) reaction (Figure 1). The number of anabolic turnovers per catabolic turnover is then determined dynamically based on the energy flux provided by the catabolic reaction and on the cost of biomass production (see Equation 1). The latter is computed accounting for biomass synthesis cost and a constant “dissipation” cost per amount of biomass, based on recent estimations [28]. Therefore, this model implements a dynamic biomass yield based on energy considerations. By lumping all the catabolic energy that is not incorporated into biomass as a constant “dissipation” term, we implicitly assume that maintenance (which is then part of dissipation) is constant. While more sophisticated dynamic representations of maintenance exist [39, 40], these approaches would add more complexity to the current model, which aims to assess how a simpler, more parsimonious thermodynamic modelling approach can capture experimental population dynamics.

The specific rate of the catabolic reaction is determined by the product of a kinetic factor (*F*_*D*_), expressing enzyme kinetics, and a thermodynamic factor (*F*_*T*_), expressing the limitations arising from thermodynamic constraints (see Figure 1 and Equation 2). *F*_*T*_ accounts for the energetic feasibility of growth-supporting pathway, as well as a minimal energy requirement (*ΔG*_*min*_). The *ΔG*_*min*_ represents the concept that cells must invest some of the energy associated with each catabolic into a metabolic driving force to run that reaction, as well as into maintaining cell viability. It is assumed that such an energy investment is pathway specific and its value can be estimated from population dynamics data, as attempted here. The resulting model is parameterised for kinetic rates using available estimates for *Dv*, *Mm*, and *Mb* (see Table S1). After this parameterisation, the only unknown parameters in the system are the *ΔG*_*min*_, which we have estimated here from the data, and some of the metabolite phase exchange constants, which we have determined experimentally. This model is developed in a generic manner allowing its expansion to include additional species and metabolic conversions. This makes it adaptable to other monocultures and natural or synthetic communities (see *Materials and Methods*).

One key feature of this generic thermodynamic model, differentiating it from previous similar models is that it implements dynamic metabolic stoichiometry through a variable yield term. The use of variable yield adjusted to close the energy balance of metabolism has indeed been advocated as a necessary feature to represent anaerobic metabolism dynamics [41, 42]. Moreover, determining the yield from physical quantities (energy gradients) reduces the amount of parameters to calibrate and thus improve the identifiability of the model’s parameters [34]. Another feature of the model is the implementation of chemical speciation in order to get a more realistic representation of the pH dynamics and the chemical concentrations during the simulations. This point, while being a rather technical one, is important especially when a dissolved species involved in a metabolic pathway has also a counterpart in the gas phase (e.g. hydrogen). In such cases, the presented model accounts for the concentration of the dissolved species in the mass action ratio of a growth-supporting metabolic pathway when determining the Gibbs free energy of that pathway.

### A thermodynamic inhibition model is required to correctly capture community metabolite dynamics

The *F*_*T*_ factor in the presented model introduces a mechanism for thermodynamic inhibition in the model, as done previously [17]. The same model without this factor could be considered as a purely ‘forward reaction kinetics model’ that considers catabolism as an irreversible process, limited only by substrate concentration [16, 22] _(see Figure 1 for illustration). We evaluate these two types of models in their ability to capture metabolite dynamics in our synthetic communities. As explained above, the model without the thermodynamic factor is nested in the presented model in the sense that it results from setting the *F*_*T*_ term to one in Equation 2. When we do so and use previously determined kinetic parameters (listed in Table S1), we can apply the resulting model without thermodynamic inhibition to the experimental data. We find that such a model is not able to explain the observed experimental results (Figure *2*). In particular, this model suggests full conversion of lactate in all culture conditions, while we do find significant lactate remaining in both *Dv* monoculture and *DvMb* co-culture. This qualitative mismatch between experiment and a non-thermodynamic inhibition model is directly a result of the structure of this model. Such a model cannot account for the low energy of lactate fermentation in the absence of sulfate, and therefore incorrectly predicts that *Dv* can consume all of the lactate. Note that this result would not change if we allow fitting of kinetic parameters in the kinetic model, as there is no mechanism in the model to allow for ‘shutting down’ of lactate consumption. The thermodynamic model, instead, allows for such a mechanism through the *F*_*T*_ term in Equation 2, and as discussed in the next section, this feature allows it to better capture the experimental data.

**Figure 2.**
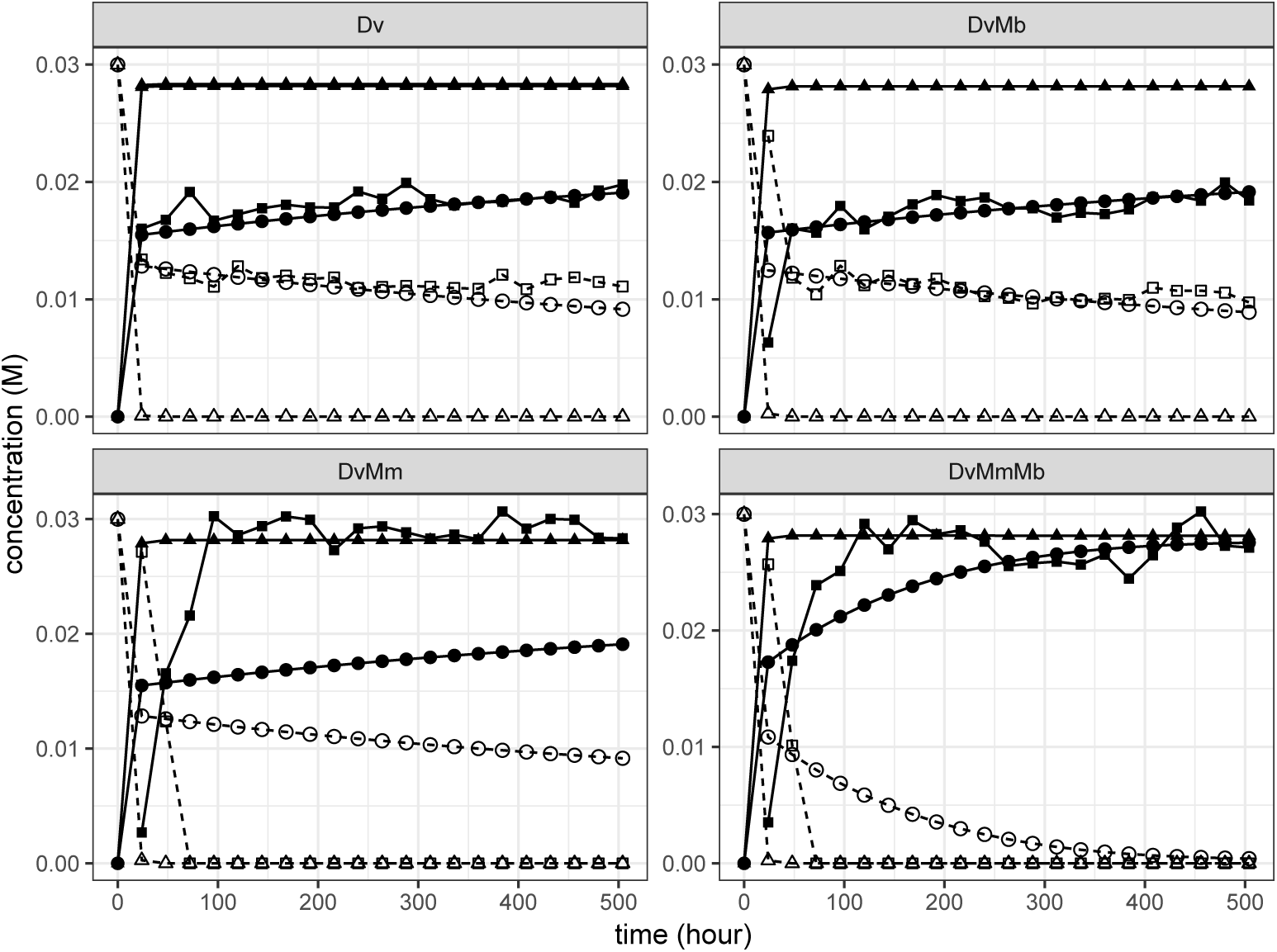
Concentration (mol/L) of acetate (solid line and filled shapes) and lactate (dashed line and empty shapes) over time (h) in the four different culture cases (*Dv*, *DvMm*, *DvMb*, *DvMmMb*) as measured in experiments (square) vs. simulated by the model (triangle or circle). Squares represent the median of the three experimental replicates, triangles represent simulations done without *F*_*T*_ factor (see Equation 2), circles represent simulations done with the *F*_*T*_ factor and using *ΔG*_*min*_ parameters obtained from calibration of the experimental data.

### Calibration of thermodynamic model allows prediction of minimal energy investments during growth with different metabolic pathways

The thermodynamic model allows better capturing of the metabolite dynamics, as shown in Figure 2. In this case, the model features additional *ΔG*_*min*_ parameters associated with the *F*_*T*_ term in Equation 2. As described above, this parameter captures the associated energy investment from each catabolic reaction into ‘running’ that reaction and into maintaining key cellular processes such as membrane potential. In order to determine this parameter for each of the possible metabolic pathways that can be used by *Dv*, *Mm* and *Mb*, we calibrated the model using an iterative fitting procedure described recently [34] (see *Materials and Methods*). The calibration process starts with an initial *ΔG*_*min*_ value of −40 kJ/mol, based on values from various sources for the Gibbs energy of formation of ATP [17, 43], and is applied using all possible combinations of the experimentally observed dynamics to result in the predicted *ΔG*_*min*_ values for each of the growth-supporting metabolic pathways (Table 2).

**Table 2:** Calibrated *ΔG*_*min*_ values for the different growth-supporting metabolic pathways modelled in this study. All values are in kJ/mol. The different rows indicate the experimental data used for the calibration. Additional results from using combinations of experimental data are provided in the Supplementary Figures S1, S2, S3 and S4.

| Culture | Calibration variable | Lactate fermentation | *Mm* Hydrogenotrophic methanogenesis | *Mb* Hydrogenotrophic methanogenesis | *Mb* Acetoclastic Methanogenesis |
| --- | --- | --- | --- | --- | --- |
| *Dv* | $\text{H}_2(\text{g})$ | -30 | | | |
|  | Ac | -25 |  |  |  |
| *DvMm* | $\text{H}_2(\text{g})$ | <-30 | >-25 | | |
|  | Ac | -15 |  |  |  |
| | $\text{CH}_4(\text{g})$ | | >-25 | | |
| *DvMb* | $\text{H}_2(\text{g})$ | -30 | | | |
|  | Ac | -25 |  |  |  |
| | $\text{CH}_4(\text{g})$ | | | >-23 | >-32 |
| *DvMmMb* | $\text{H}_2(\text{g})$ | <-31 | <-20 | >-20 | |
|  | Ac | -15 |  |  |  |
| | $\text{CH}_4(\text{g})$ | | <-30 | >-25 | >-25 |

After a set of parameters was determined by the calibration procedure for each combination of observed variables, a parametric sweep was performed to determine whether the obtained values correspond to an optimum that minimizes the distance between simulation and observation. We assess this by plotting an error function (see *Materials and Methods*) for each calibrated parameter value (Figure 3 and S1-S4). The shape of the error function around the calibrated values of each parameters indicates that the *ΔG*_*min*_ of the lactate fermentation pathway has a clear optimum regarding the output variables considered (acetate and H_2_ in gas phase), and lies between −30 and −15kJ/mol. The *ΔG*_*min*_ of the hydrogenic and acetoclastic methanogenesis pathways cannot be given an exact estimate but rather boundaries, between 0 and −20kJ/mol for hydrogenic methanogenesis and less negative than −40kJ/mol for hydrogenic methanogenesis (Figure 3A). The *ΔG*_*min*_ parameters for *Dv*’s sulfate respiration pathways could not be calibrated with the present experimental data, presumably because sulfate respiration occurs relatively quickly compared to the time-resolution of the available experimental data.

**Figure 3.**
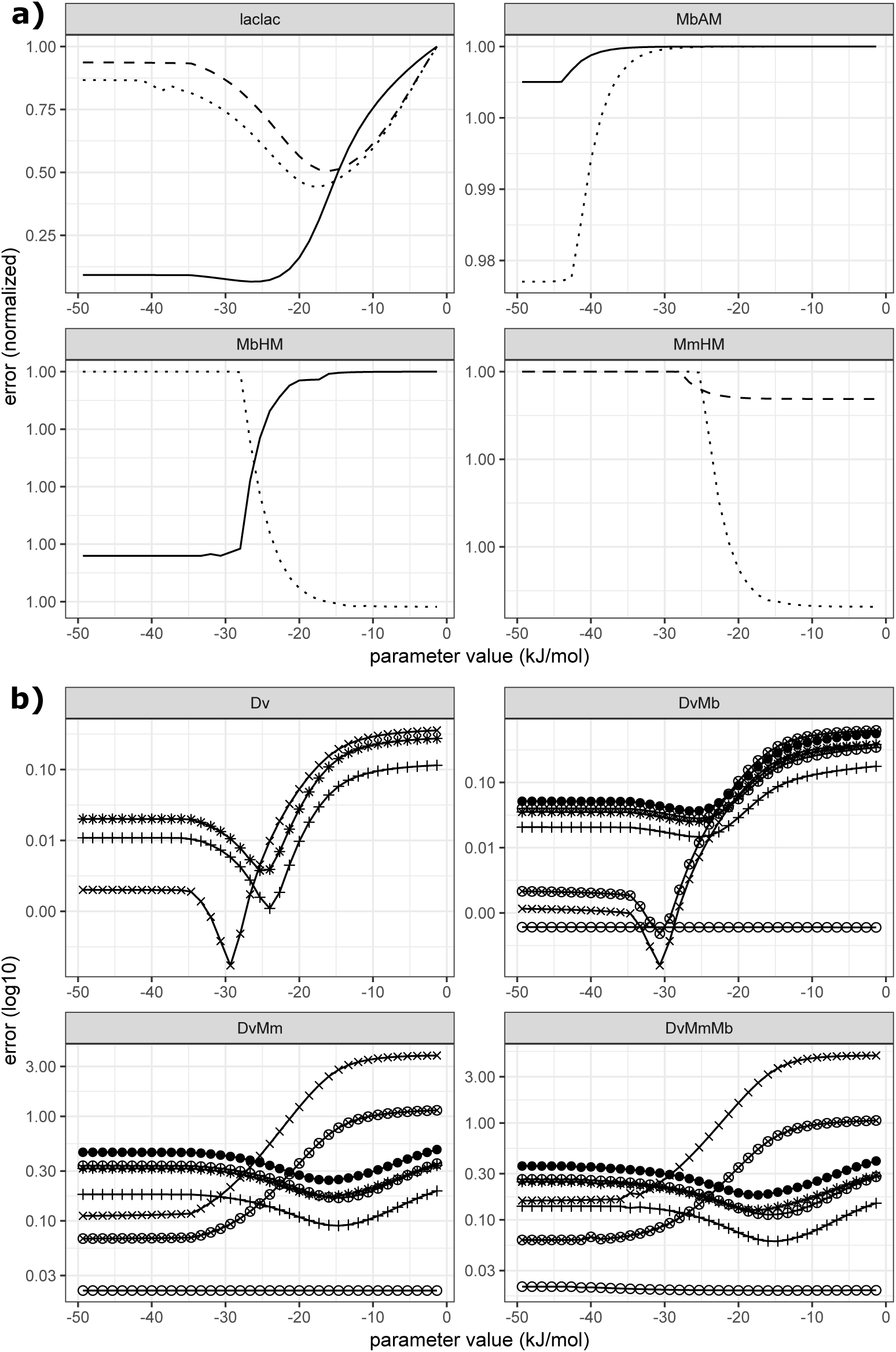
Sum of squared differences (error) between the experimentally observed variable(s) and the model prediction (y-axis) as a function of the value of the *ΔG*_*min*_ of various pathways (x-axis). **(A)** Normalised error between experimentally observed and predicted concentration of CH_4_(g) and H_2_(g) over time, depending on the values of *ΔG*_*min*_ for various pathways; lactate fermentation by *Dv* (laclac), acetoclastic methanogenesis by *Mb* (AM), hydrogenotrophic methanogenesis by *Mb* (HM) and hydrogenotrophic methanogenesis by *Mm* (HM). The error results shown are for the case using the experimentally observed H_2_(g) and acetate concentrations as the comparison. Results for different cultures are indicated with the line properties; solid line for *DvMb*, dashed line for *DvMm*, dotted line for *DvMmMb*. **(B)** Log of the error, as a function of the estimated value of the *ΔG*_*min*_ of the lactate fermentation pathway. Each tile corresponds to the results from a different culture case. The experimentally observed variables on which the error is computed are; acetate (plus), H_2_(g) (cross), CH_4_(g) (circle), acetate and H_2_(g)-(plus and cross), H_2_(g) and CH_4_(g) (circle and plus), acetate and CH_4_(g) (circle and cross), and H_2_(g), acetate and CH_4_(g) (black filled circles).

As far as we are aware, estimation of the *ΔG*_*min*_ parameter has only been attempted before in few studies [20, 30]. The minimal free energy for the three metabolic pathways of *Dv* (lactate fermentation and sulfate respiration with lactate or hydrogen) has been calibrated against experimental data only from monocultures grown in the presence of sulfate, and using a model similar to that presented here [30]. The experimental design in that study was different, using solely a monoculture (rather than monoculture and communities as we do here), sampling at shorter time intervals and using higher sulfate concentration than this study. Perhaps due to such differences, the estimated value from that study for lactate fermentation was −39.5 kJ/mol, relatively higher (i.e. more energy investment required as driving force and into maintenance) than found here. Possibly due to its use of higher sulfate concentration and shorter sampling intervals, that study was able to estimate the *ΔG*_*min*_ for both lactate and H_2_ respiration on sulfate, as −44.66 kJ/mol. It should be noted that the model utilised by that study is different from the model used here, in that it uses gas partial pressure when calculating reaction free energies and has taken a static approach to model biomass yield. A theoretical study aimed at estimating energetic parameters for several metabolic pathways, including methanogenesis, using existing data [44], but, it used a notion of minimum energy threshold that requires assumptions about the underlying metabolic reactions. The final minimum energy threshold is then expressed in terms of ATP molecules produced per metabolic pathway turnover. The resulting predictions from that study cannot be directly translated into a minimum energy threshold if we assume that the Gibbs free energy carried by an ATP molecule varies dynamically with the state of the cell’s ATP pool, and therefore cannot be compared directly to the results presented here.

It is interesting to note that when a parameter sweep shows the existence of an optimal value or range, these depend on the experimental variable and culture used for calibration (see Table 2, Figure 3B). There are two possible explanations for this observation. Firstly, there may be metabolic pathways being catalysed by the populations in the different experimental batches that are not represented by the model. If such pathways involve a specific metabolite, then calibrations performed on that metabolite vs. some other metabolite might differ. Such an explanation, while theoretically possible, does not fit with the fact that the presented model accounts for all key pathways known to be catalysed by *Dv*, *Mm*, and *Mb*. The second possible explanation is that the minimum energy threshold of a given metabolic pathway depends on the experimental conditions. Indeed, the concept of a minimum energy threshold for growth aims to conceptualise energy invested as metabolic driving force as well as cellular maintenance. Both these energetic investments are expected to be a function of culture and cellular conditions, including specific cellular details such as Mg^+2^ concentration [45]). The minimum energy threshold of a growth-supporting pathway is then expected to be dynamic. However, accurately predicting those dynamic variations would require to implement a detailed model of the populations’ metabolic networks and cellular states.

## CONCLUSIONS

Here, we have developed and presented a generic thermodynamic model for capturing population and metabolite dynamics in a microbial community. The model implements specific features that have been proposed and advocated over the last two decades [22, 28, 33, 41] by introducing factors based on first principles (thermodynamic limitation of reaction rate [47]). As such, it overcomes the limitations of modelling approaches solely based on Michaelis-Menten-type kinetics and empirically calibrated product inhibitions (such as with the ADM1 model [13]). We applied this model to capture the dynamics of synthetic communities composed of a sulfate reducer and two methanogens. We also used the model to attempt an estimation of the minimum energy thresholds of the different growth-supporting metabolic pathways found in these organisms, sulfate respiration, lactate fermentation, and hydrogenotrophic and acetoclastic methanogenesis. Our findings show that the presented model, while simple, is indeed able to capture some of the thermodynamic limitations occurring in the observed dynamics. Further, the use of the model on experimental data allows for the prediction of the minimum energy requirements for sulfate fermentation and hydrogenotrophic and acetoclastic methanogenesis.

Our calibration results also shed light on the limitations of the thermodynamic approach employed. In particular, despite improvements over a non-thermodynamic model, the presented model was not able to fully capture experimental metabolite and population dynamics and its combination with experimental data did not allow to precisely determine the minimal free energies for all modelled pathways. These limitations might be inherent in the structure of the model or in the experimental design used here, or a combination of the two. The latter can be addressed particularly by collecting more and higher resolution temporal data from similar experimental systems. The former will probably require increasing the complexity of the presented model. In particular, considering minimum energy thresholds as a constant feature of the system may be too simplistic and might instead require including elements of metabolic pathway dynamics within the bacterial growth models.

Thermodynamic constraints that we have endeavoured to predict here are one of the few phenomena that can be safely assumed to apply for all growth-supporting metabolic pathways. A sound basis for the description of this fundamental constraint applying to metabolic dynamics is thus necessary before attempting to assess and calibrate the extent of higher order phenomena such as genetic regulation or resource allocation [46, 47]. While further dedicated experiments and more complex models are necessary to improve the accuracy of dynamics predictions, the presented work provides a step towards this aim. The presented model expands previous efforts of minimal energy estimates from monocultures [30] and combines several recently proposed model features such as dynamic growth yield [22, 41] with additional features such as modelling of multiple pathways within individual species, phase exchange dynamics, and pH. As such, its further use and assessment will facilitate thermodynamic modelling of microbial community dynamics and estimation of energetic parameters, helping the development of more predictive microbial community dynamics models.

## ACKNOWLEDGEMENTS

We thank Dr. Fred Farrell for his contributions to the software development of an earlier version of the presented model.

## AUTHOR’S CONTRIBUTIONS

O.S.S. designed the overall study and contributed to model development. J.C contributed to design of experiments and performed them. M.W contributed to the design of the model and its initial implementation. H.D designed the model, implemented it and performed data fitting and model analyses. All authors contributed to the writing of the manuscript and have given approval to the final version.

