## Supplementary Material for "Thermodynamic modelling of synthetic communities predicts minimum free energy requirements for sulfate reduction and methanogenesis"

### Supplementary material
### for

**Thermodynamic modelling of synthetic communities predicts minimum free energy requirements for sulfate reduction and methanogenesis.**

Hadrien Delattre, Jing Chen, Matthew Wade, Orkun S Soyer

This supplementary text provides additional Tables and Figures as referenced from the main text.

| Organism | Metabolism | Parameter | Value | Citation |
|---|---|---|---|---|
| *Dv* | Lactate fermentation | $v_{max}$ | 1.03 | Noguera |
| *Dv* | Lactate respiration | $v_{max}$ | 0.942 | Noguera |
| *Dv* | $H_2$ respiration | $v_{max}$ | 0.848 | Noguera |
| *Mm* | Hydrogenotrophic methanogenesis | $v_{max}$ | 8.49e-4 | Robinson & Tiedje |
| *Mm* | Hydrogenotrophic methanogenesis | $K_m$ ($H_{2,aq}$) | 3.75e-6 | Robinson & Tiedje |
| *Mb* | Acetoclastic methanogenesis | $v_{max}$ | 2.68e-4 | Westermann, Ahring & Mah |
| *Mb* | Acetoclastic methanogenesis | $K_m$ (Acetate) | 4.5e-3 | Westermann, Ahring & Mah |
| *Mb* | Hydrogenotrophic methanogenesis | $v_{max}$ | 1.17e-3 | Westermann, Ahring & Mah |
| *Mb* | Hydrogenotrophic methanogenesis | $K_m$ ($H_{2,aq}$) | 9.5e-6 | Westermann, Ahring & Mah |

**Table S1:** Value and literature source for the parameters of the enzymatic kinetics of the modelled pathways

| Chemical | Henry's Constant at 298K (mol/(m$^3$.Pa)) | $\Delta H_{sol}/R$ (K) | $k_La$ (1/h) | Citation |
|---|---|---|---|---|
| $CO_2$ | 3.4e-4 | 2400 | 1.10e-1 | This study |
| $H_2$ | 7.8e-6 | 500 | 1.33e-1 | This study |
| $CH_4$ | 1.43e-5 | 1600 | 0 | This study |
| $H_2S$ | 1.0e-3 | 2100 | 8.33e-2 | (Noguera *et al.* 1998) |
| $NH_3$ | 5.9e-1 | 4200 | 1.21e-1 | Estimated as geometric mean between $CO_2$ and $H_2$ $k_La$ |

**Table S2**: Henry's constants and gas/liquid transfer rate parameters used for the simulations.

| Chemical | pK constant(s) |
|---|---|
| $C_2H_4O_2 \rightleftharpoons C_2H_3O_2^-$ | 4.38 |
| $C_3H_6O_3 \rightleftharpoons C_3H_5O_3^-$ | 3.78 |
| $H_3PO_4 \rightleftharpoons H_2PO_4^- \rightleftharpoons HPO_4^{-2} \rightleftharpoons PO_4^{-3}$ | 1.79; 6.95; 12.89 |
| $CO_2(aq) \rightleftharpoons HCO_3^- \rightleftharpoons CO_3^{-2}$ | 6.35; 10.33 |

| | |
|---|---|
| $H_2S(aq) \rightleftharpoons HS^-$ | 7.00 |
| $HSO_4^- \rightleftharpoons SO_4^{-2}$ | 1.99 |
| $NH_4^+ \rightleftharpoons NH_3(aq)$ | 9.25 |

**Table S3:** Acid/base equilibria considered in the simulations and their respective pK constant (at 310.15 K)

| **Case** | **OD600 measurement at day 0** | | | **Average OD600** | **Biomass concentration estimate (C-mol/L)** |
|---|---|---|---|---|---|
| | **Replicate 1** | **Replicate 2** | **Replicate 3** | | |
| ***Dv*** | 0.379 | 0.375 | 0.411 | 0.388 | 2.02e-3 |
| ***DvMm*** | 0.028 | 0.027 | 0.021 | 0.025 | 3.15e-4 |
| ***DvMb*** | 0.110 | 0.115 | 0.158 | 0.128 | 5.34e-4 |
| ***DvMmMb*** | 0.05 | 0.038 | 0.053 | 0.047 | 5.53e-4 |

**Table S4:** Initial biomass concentration in each experimental case

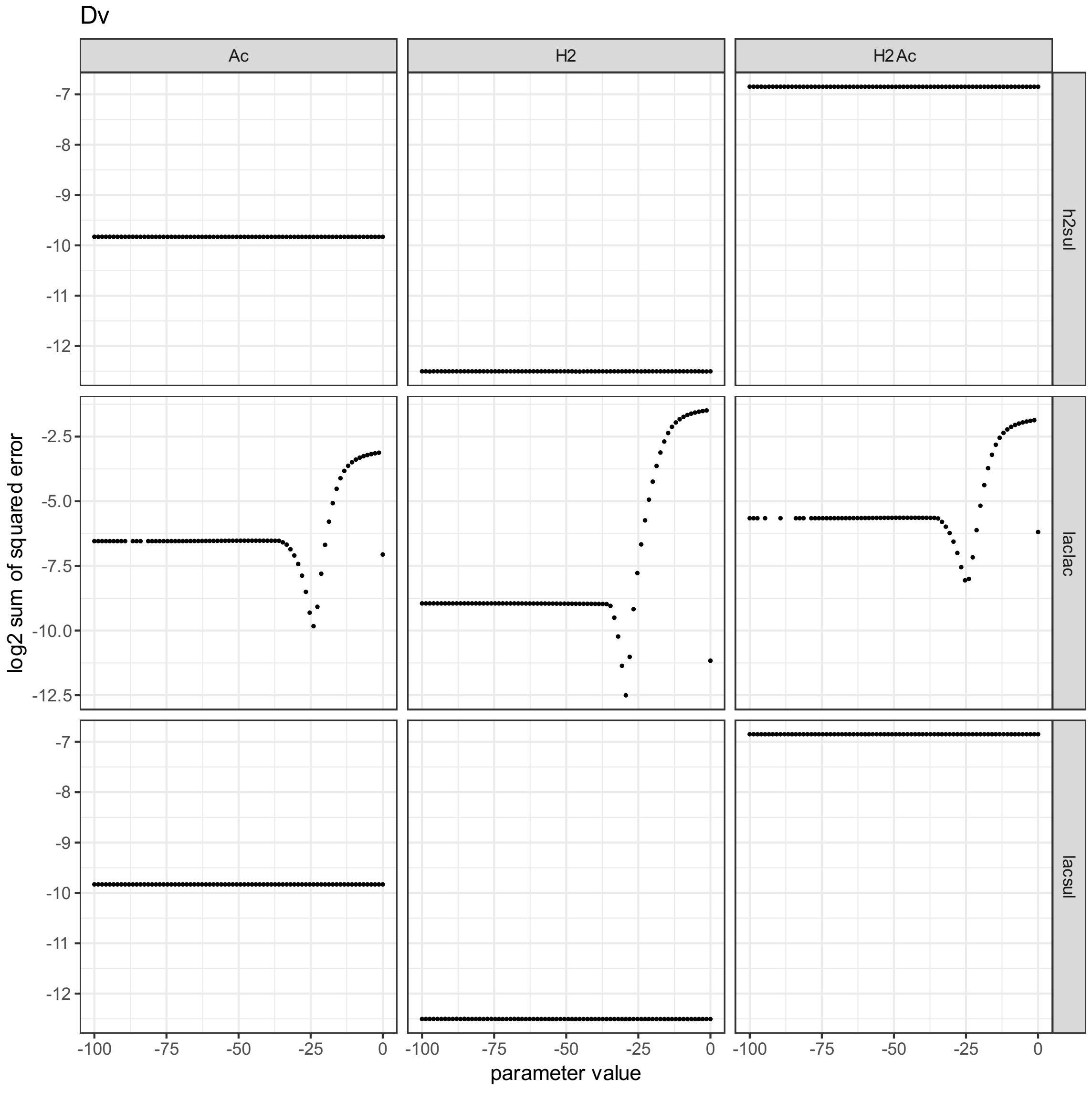


**Figure S1:** Sum of squared differences (error) between the experimentally observed variable(s) and the model prediction (y-axis of each tile) of the *Dv* monoculture, as a function of the value

of the $\Delta G_{min}$ parameter of various pathways (x-axis of each tile), depending on the calibration variable (x-axis of the figure) and the $\Delta G_{min}$ parameter considered (y-axis of the figure; laclac: lactate fermentation, lacsul: lactate respiration on sulfate, h2sul: hydrogen respiration on sulfate).

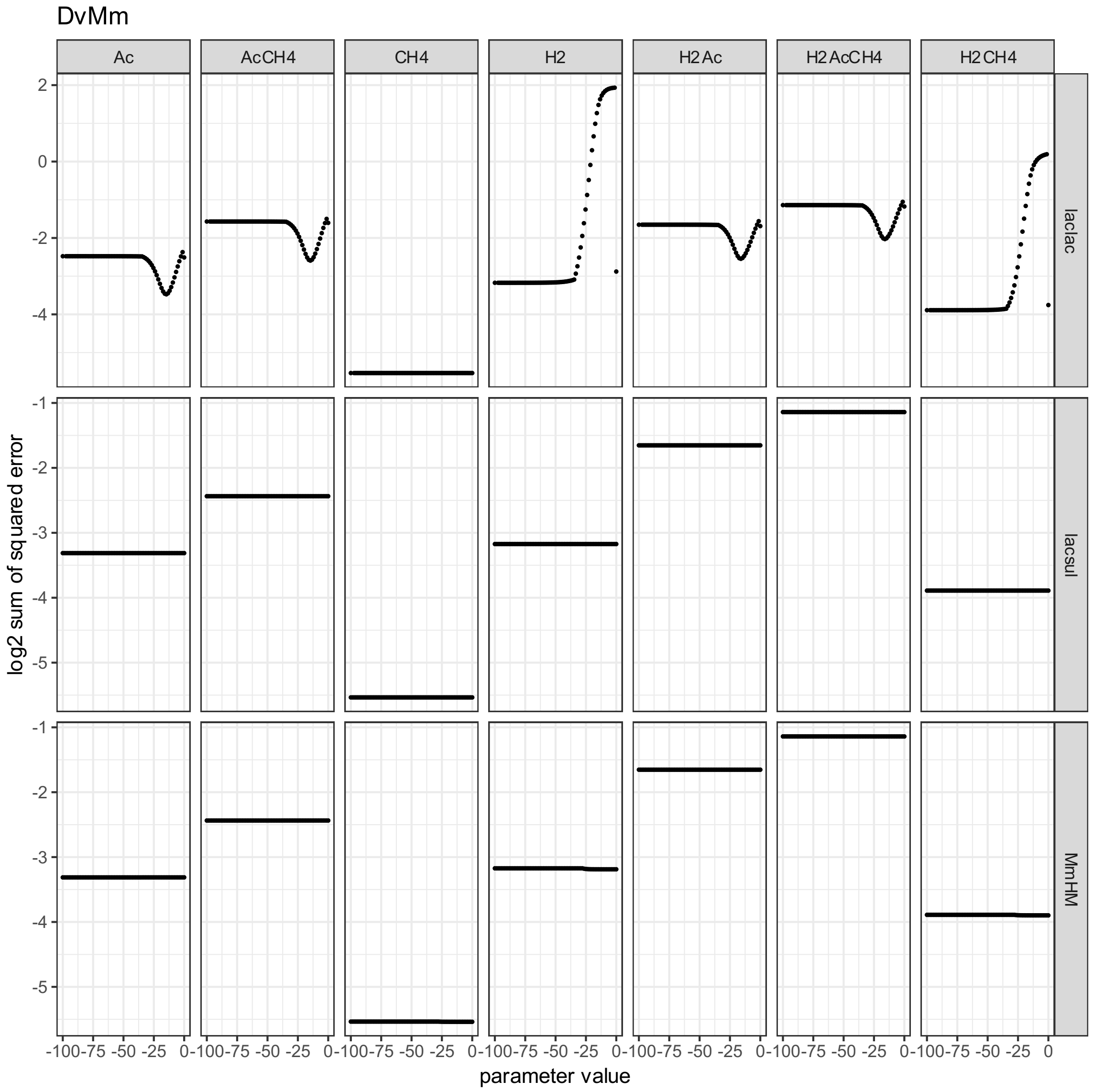


**Figure S2:** Sum of squared differences (error) between the experimentally observed variable(s) and the model prediction (y-axis of each tile) of the *DvMb* coculture, as a function of the value of the $\Delta G_{min}$ parameter of various pathways (x-axis of each tile), depending on the calibration variable (x-axis of the figure) and the $\Delta G_{min}$ parameter considered (y-axis of the figure; laclac: lactate fermentation by *Dv*, lacsul: lactate respiration on sulfate by *Dv*, MbAM: acetoclastic methanogenesis by *Mb*, MbHM: hydrogenotrophic methanogenesis by *Mb*).

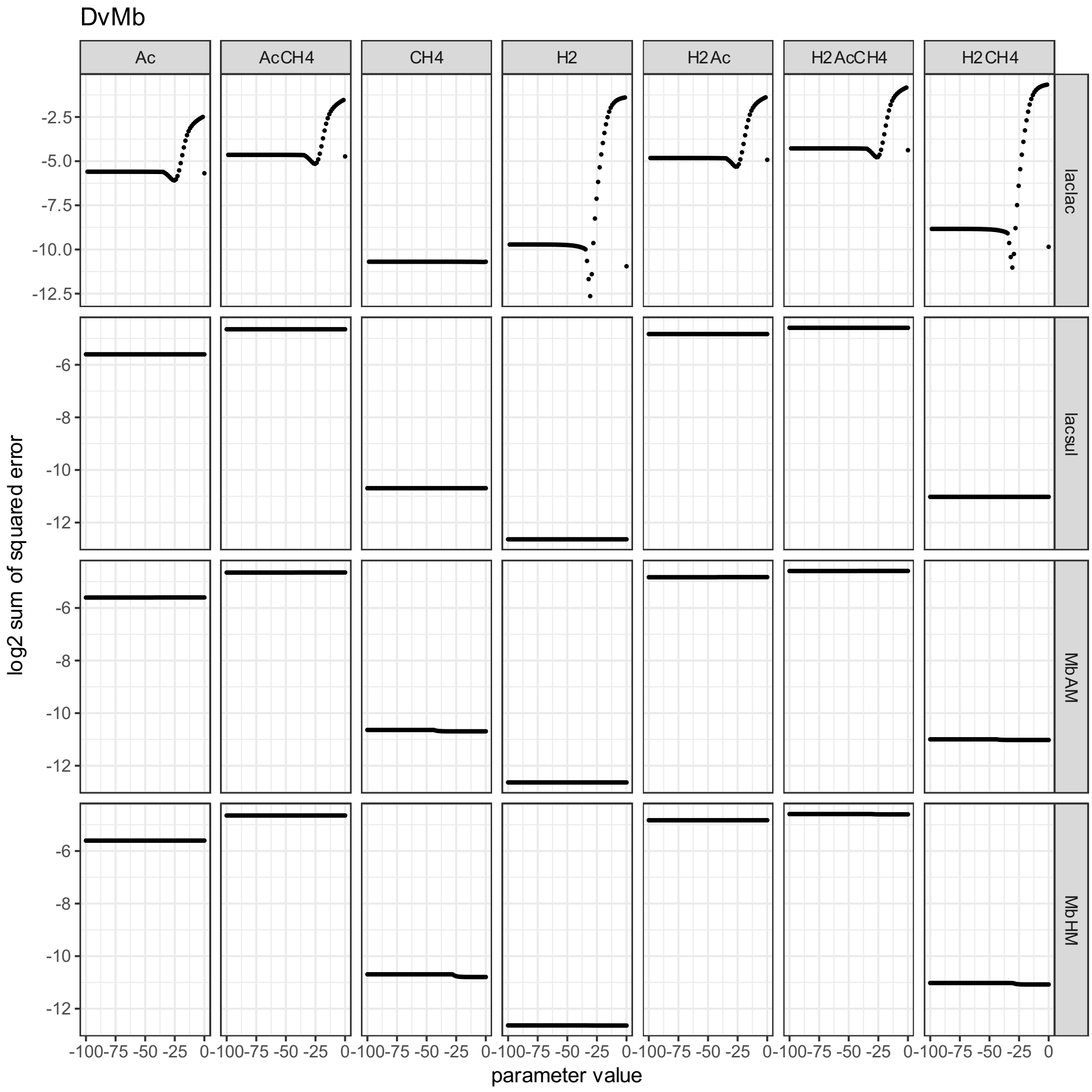


**Figure S3:** Sum of squared differences (error) between the experimentally observed variable(s) and the model prediction (y-axis of each tile) of the *DvMm* coculture, as a function of the value of the $\Delta G_{min}$ parameter of various pathways (x-axis of each tile), depending on the calibration variable (x-axis of the figure) and the $\Delta G_{min}$ parameter considered (y-axis of the figure; laclac: lactate fermentation by *Dv*, lacsul: lactate respiration on sulfate by *Dv*, MmHM: hydrogenotrophic methanogenesis by *Mm*).

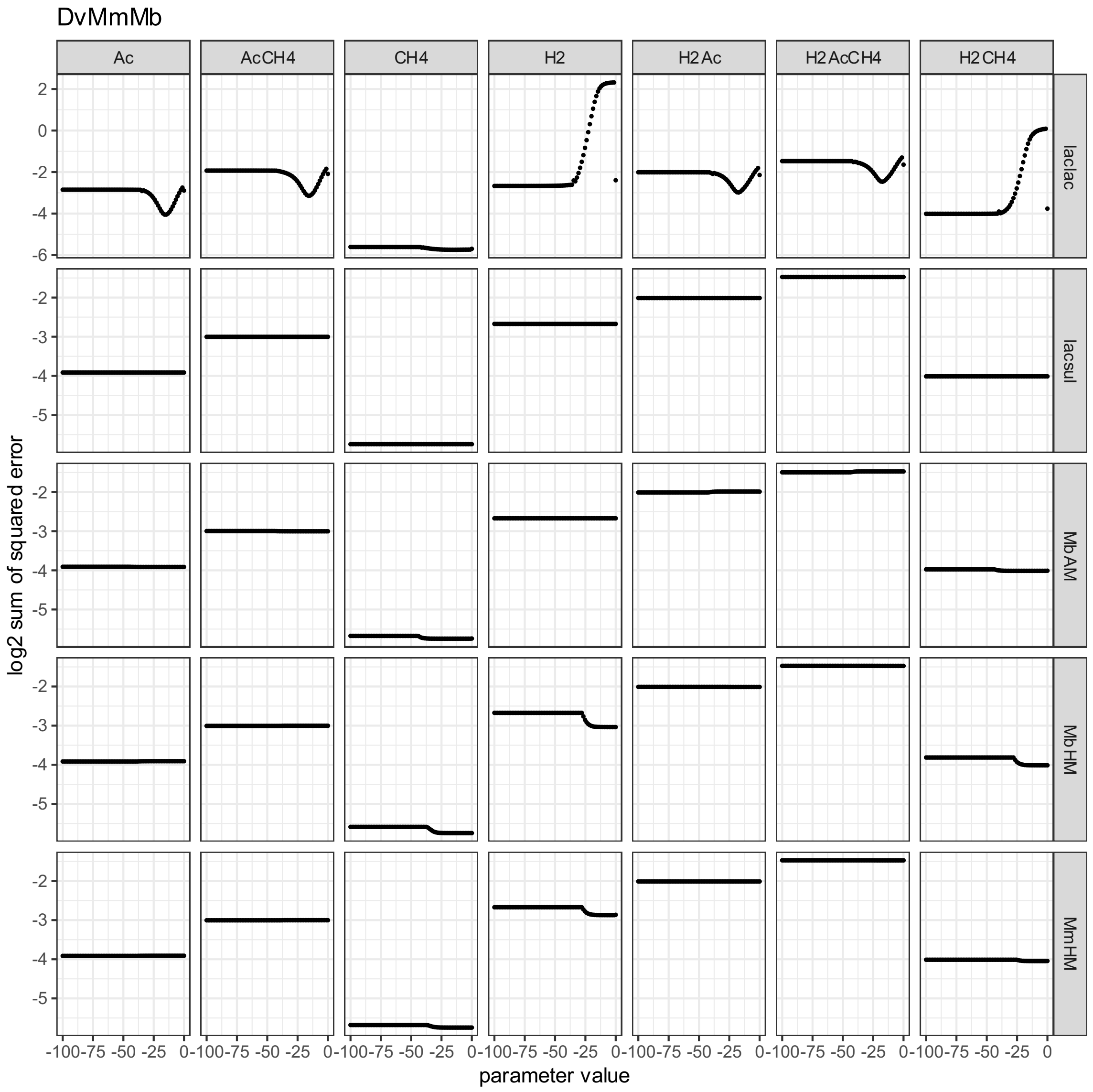


**Figure S4:** Sum of squared differences (error) between the experimentally observed variable(s) and the model prediction (y-axis of each tile) of the *DvMmMb* coculture, as a function of the value of the $\Delta G_{min}$ parameter of various pathways (x-axis of each tile), depending on the calibration variable (x-axis of the figure) and the $\Delta G_{min}$ parameter considered (y-axis of the figure; laclac: lactate fermentation by *Dv*, lacsul: lactate respiration on sulfate by *Dv*, MbAM: acetoclastic methanogenesis by *Mb*, MbHM: hydrogenotrophic methanogenesis by *Mb*, MmHM: hydrogenotrophic methanogenesis by *Mm*).
